# Sex- and hepatocyte PPARγ-dependent effects of an obesogenic dietary approach to induce MASH with fibrosis in mice

**DOI:** 10.64898/2026.02.25.707976

**Authors:** Marta Sierra-Cruz, Izabela Hawro, Samuel M. Lee, Jose Muratalla, Jose Cordoba-Chacon

**Affiliations:** Department of Medicine. Division of Endocrinology, Diabetes and Metabolism. University of Illinois at Chicago, Chicago. IL

**Author notes:** **Corresponding author’s contact Information**: Jose Cordoba-Chacon, PhD. Department of Medicine, Section of Endocrinology, Diabetes and Metabolism. 835 S. Wolcott Ave (North Entrance) Suite E625. M/C 640. Chicago, IL. MSC, IH, SML are co-first authors of this study.

## Abstract

Mouse models of metabolic dysfunction-associated steatotic liver disease (MASLD) are valuable tools for identifying novel molecular mechanisms that drive progression from MASLD to metabolic dysfunction-associated steatohepatitis (MASH). However, generating a clinically relevant MASLD/MASH mouse model with obesity and peripheral metabolic dysfunction remains a challenge. In this study, we fed two different MASH-inducing diets to male mice with pre-existing high-fat (HF) diet-induced obesity. While a HF diet containing 40% Kcal from fat (mostly corn-oil shortening), 2% cholesterol, and 22% fructose reduced adiposity in these mice, a high-fat diet with 60% Kcal from fat (mostly lard), containing 2% cholesterol and supplemented with 10% fructose in the drinking water (HFC+Fr diet) promoted body weight and fat mass gain. Of note, 24 weeks of the HFC+Fr diet induced obesity, metabolic dysfunction, and liver steatosis in male and female mice, and promoted MASH with fibrosis in male mice. Furthermore, the HFC+Fr diet increased the expression of hepatocyte peroxisome proliferator-activated receptor γ (*Pparg*), but the knockout of *Pparg* in hepatocytes (*Pparg*^ΔHep^) reduced the development of MASH and fibrosis in male mice. In addition, the expression of key hepatic genes involved in methionine metabolism was downregulated by the HFC+Fr diet and upregulated by *Pparg*^ΔHep^ only in male mice. Overall, the HFC+Fr diet is obesogenic and promotes MASLD in both male and female mice. However, the HFC+Fr diet promotes MASH in a sex- and hepatocyte *Pparg*-specific manner, which may be associated with downregulation of hepatic methionine metabolism.

**New & Noteworthy:** We explored how a new dietary intervention with fructose in the drinking water and added cholesterol to a high-fat diet extensively used to induce obesity and insulin resistance, promotes the onset of MASLD with obesity and metabolic dysfunction in male and female mice. This clinically relevant model of MASLD shows increased expression of hepatocyte PPARγ in both male and female mice, but only male mice have PPARγ-dependent impaired methionine metabolism and develop MASH with fibrosis.

## Introduction

Metabolic dysfunction-associated steatotic liver disease (MASLD) is currently defined as the accumulation of fat in the liver (steatosis) accompanied by at least one cardiometabolic risk factor without other identifiable secondary causes of steatosis (1). MASLD affects to 38% of adults in the United States, and roughly 25% of patients with MASLD develop metabolic dysfunction-associated steatohepatitis (MASH) (2). MASH is characterized by hepatocellular ballooning, lobular inflammation, and liver fibrosis, which increases the risk of progressing to cirrhosis which can then develop into hepatocellular carcinoma (3,4). Over recent decades, researchers have used different mouse models to study the molecular and cell-specific mechanisms that drive the progression from MASLD to MASH (5,6). Some of these models do not fully reproduce the most common metabolic abnormalities in patients with MASLD and MASH, because nutrient-deficient diets that promote MASH increase insulin sensitivity and reduce adiposity (7,8). However, some Western diets and high-fat (HF) diets that promote MASLD are reliable tools to induce human-like MASLD/MASH in mice (9,10). Nonetheless, the composition of these special diets may not fully replicate the full spectrum of metabolic abnormalities associated with human steatohepatitis (5,11).

Our group has investigated the role of hepatocyte peroxisome proliferator-activated receptor gamma (*Pparg,* PPARγ) in the development of MASLD (12) using mice that were fed the classic HF diets that induce obesity, insulin resistance, and liver steatosis (13,14), the methionine and choline-deficient (MCD) diet that induces steatohepatitis (15), or a high-fat, cholesterol, and fructose (HFCF) diet that induces MASH without severe metabolic dysfunction (11,16). The use of hepatocyte-specific *Pparg* knockout (*Pparg*^ΔHep^) mice in these studies indicates that hepatocyte *Pparg* expression is associated with HF-mediated liver steatosis, MCD-mediated fibrosis, and HFCF-mediated inflammation and fibrosis (12). It should be noted that the nutrient-specific composition of the HFCF diet we previously used prevents the induction of a clear baseline for obesity and insulin resistance (11,16). Thus, in this study, we compared our previous HFCF diet with an alternative diet to generate a clinically relevant model of MASLD/MASH with severe obesity and insulin resistance in both control and *Pparg*^ΔHep^ mice. More specifically, we have assessed the early effects of two MASH-inducing diets on body weight gain, adiposity, and MASH in male mice with pre-existing diet-induced obesity and reproduced the effects that *Pparg*^ΔHep^ has on liver histology in mice with diet-induced MASH. Furthermore, we assessed whether the nutrient composition of a new dietary intervention induces MASH, obesity, and insulin resistance in male and female mice to generate a clinically relevant model of MASH. Overall, this study shows that the HFC+Fr diet promotes MASH with fibrosis in male mice with pre-existing diet-induced obesity in a hepatocyte *Pparg*-dependent manner without reducing adiposity. Furthermore, this new dietary intervention promotes diet-induced obesity and liver steatosis in both male and female mice. However, only male mice develop MASH with fibrosis in a hepatocyte *Pparg*-dependent manner.

## Material and Methods

### Mouse models

The studies with mice were approved by the Institutional Animal Care and Use Committee of the University of Illinois Chicago. *Pparg* floxed mice (17) in a C57BL/6J background were purchased from Jackson Laboratories (Strain 004584, B3.129-Ppargtm2Rev/J, Bar Harbor, ME), and bred as homozygotes under controlled temperature (22-24°C) and humidity in a specific-pathogen-free facility with 14 h light/10 h dark cycle (lights on at 6:00 am).

To induce MASH in mice with pre-established diet-induced obesity, we generated two cohorts of five to seven-week-old chow-fed *Pparg* floxed littermate male mice. *Pparg* floxed mice were fed a high-fat diet containing 60% kcal from fat (HF or HFD, Cat #D12492, Research Diets, Inc) for 16 weeks or a nutrient- and fiber source-matched low-fat diet containing 10% kcal from fat (LF or LFD, Cat #D12450J, Research Diets, Inc, New Brunswick, NJ, Supplementary Table 1). After 16 weeks of HF feeding, a subset of HF-fed *Pparg* floxed mice were injected in the lateral tail vein with a single bolus of 1.5 × 10¹¹ genome copies/mouse adeno-associated virus encoding Cre recombinase under a hepatocyte-specific promoter (thyroxin binding globulin, TBG, AAV8-TBG-Cre, #107787-AAV8, Addgene, Watertown, MA) to generate *Pparg*^ΔHep^ mice, and another subset of *Pparg* floxed mice fed a HF diet was injected with AAV-TBG-Null (#105536-AAV8, Addgene) to generate control mice, as we have previously published (14). Two weeks after AAV injections in the cohort #1, half of the HF-fed control or *Pparg*^ΔHep^ mice were switched to a high-fat, cholesterol and fructose diet (HFCF, 40% kcal fat from hydrogenated corn oil, 2% cholesterol and 22% fructose; Cat #D16010101, Research Diets, Inc, Supplementary Table 1). One week after AAV injections in the cohort #2, half of HF-fed control or *Pparg*^ΔHep^ mice were switched to a high-fat and cholesterol diet (HFC, 60% kcal from lard fat and 2% cholesterol; Cat# D12120101, Research Diets, Inc, Supplementary Table 1) supplemented with 10% fructose in the drinking water (HFC+Fr). The rest of the HF-fed control or *Pparg*^ΔHep^ mice remained on the HF diet. A group of littermate *Pparg* floxed mice fed a LF diet was simultaneously injected with AAV8-TBG-Null and served as baseline controls.

We used additional cohorts of mice to assess the effects of HFC+Fr diet on diet-induced obesity and MASH in male and female mice (Cohort #3) and on metabolic rate of male mice (Cohort #4). For the cohort #3 we used nine to eleven-week-old chow-fed *Pparg* floxed littermate male and female mice, and for the cohort #4 we used eleven to fourteen-week-old chow-fed *Pparg* floxed littermate male mice. These mice were treated with AAV vectors to generate control or *Pparg*^ΔHep^ mice as indicated above. Two weeks later, a subset of control and the *Pparg*^ΔHep^ mice were switched to an HFC+Fr diet for 24 weeks. A subset of control mice was fed a LF diet.

After 25-26 weeks (Cohort #1 and #2), or 24 weeks (Cohort #3) of special diets, mice were euthanized by decapitation after 4h-food withdrawal at 8:00 am. Trunk blood was collected with EDTA-coated BD Microtainer® (BD, Franklin Lakes, NJ), and plasma was obtained after centrifugation and stored at -20°C. Livers were removed, weighed, and fixed in formalin or snap-frozen in liquid nitrogen and stored at -80°C. White adipose tissue: urogenital, mesenteric, retroperitoneal, and subcutaneous fat-depots, as well as brown adipose tissue were collected and weighed.

### Whole body composition and metabolic endpoints

The effect of diet-induced MASH on whole-body composition (fat, lean, and free fluid mass) was assessed with a minispec LF50 body composition analyzer (Bruker, Billerica, MA) in Cohorts #1-3. In cohort #3, a glucose tolerance test (GTT) was performed after 21 weeks of special diets as previously reported (16). In cohort #4, energy expenditure, oxygen consumption, carbon dioxide output, respiratory exchange ratio, and food consumption were measured using Promethion Systems (Sable Systems International, Las Vegas, NV) for four consecutive days. Plasma levels of non-esterified fatty acid (NEFA), triglycerides (TG), cholesterol (Wako Diagnostics, Richmond, VA), and alanine aminotransferase (ALT, Pointe Scientific, Canton, MI) were measured with colorimetric assays. Plasma insulin levels were measured with a commercial ELISA (Mercodia, Winston-Salem, NC) and blood glucose was measured with a commercial human glucometer (Roche, Indianapolis, IN). For hepatic TG and cholesterol quantification, livers were processed as previously reported (18).

### Liver histology

Formalin-fixed livers were processed by the Research Histology Core of the University of Illinois Chicago. Paraffin-embedded 5 µm sections were stained with hematoxylin-eosin (H&E) to determine the MASLD activity score in a blind manner following the criteria described by *Kleiner et al.* (19). A MASLD activity score greater than 5 was considered MASH. Moreover, liver tissue sections were stained with picrosirius red/fast green (Cohort #1 and #2) or picrosirius red (Cohort #3) to quantify the area of fibrosis using a specific macro with *ImageJ* as previously described (16)

### Hepatic gene expression: quantitative PCR analysis, RNAseq, and western-blot

Total RNA was extracted using Invitrogen™ TRIzol™ Reagent (Thermo Fisher Scientific, Carlsbad, CA), and treated with RQ1 RNase-Free DNase (Promega, Madison, WI). DNA-free RNA was reverse-transcribed into cDNA using the First Strand cDNA Synthesis Kit (Thermo Scientific, Waltham, MA), and gene expression was quantified with quantitative real-time PCR using Brilliant III Ultra-Fast QPCR Master Mix (Agilent Technologies, Santa Clara, CA) and gene-specific primers listed in Supplementary Table 2 (20,21). Liver RNA obtained from Cohort #3 was used for RNAseq. Libraries preparation, sequencing, and bioinformatics analysis of RNA-seq were performed by Novogene (Novogene, Inc, Sacramento, CA), as previously described (16). The raw and processed files, including those with the differentially expressed genes (DEGs), have been deposited and published in the Gene Expression Omnibus (GEO) under accession number #GSE315007. Proteins were extracted from frozen livers with RIPA lysis buffer supplemented with phosphatases and proteases inhibitors (Milipore Sigma, Burlington, MA) using Red Rino bead lysis kit and the bullet blender homogenizer (Next Advance, Troy, NY). Isolated proteins were quantified using Bicinchoninic acid (BCA) and separated in 26-well Criterion TGX gels (Bio-Rad, Hercules, CA) and transferred onto nitrocellulose membranes as previously described (13,16). Ponceau S staining was performed to confirm protein transfer and used for protein normalization (22). Then, ponceau S stain was removed, and the membranes were incubated in the blocking buffer: 5% non-fat dry milk in Tris-buffered saline with 0.1% Tween-20 (TBST) for one hour at room temperature, and incubated at 4°C overnight with either PPARγ (1:500, C26H12, Cell Signaling Technologies, Danvers, MA, USA), COL1A1 (1:1000, E8F4L, Cell Signaling Technologies,), CIDEC (1:500, NB100-430,Novus Biologicals, Centennial, CO), BHMT (1:10000, ab96415, Abcam, Waltham, MA). Goat anti-rabbit immunoglobulin G (H+L)-horseradish peroxidase conjugate (1:2000 for 2 h [PPARγ and CIDEC], 1:10000 for 1h [COL1A1], and 1:20000 for 1h [BHMT], Bio-Rad) was used as secondary antibody, and the immunoreactive specific signals were detected with a ChemiDoc Imager and quantified with ImageLab software (Bio-Rad).

### Statistical analysis

Data were analyzed with GraphPad Prism (Graphpad, San Diego, CA). In the cohorts #1 and #2, we used Student’s *t test* to assess the effect of HF diet on control mice, and two-way analysis of variance (ANOVA) with a post-hoc Tukey’s HSD multiple comparisons test to assess the effect of HFCF or HFC+Fr diets, and *Pparg*^ΔHep^. In the cohort #3, we used Student’s *t- test* to assess the effect of HFC+Fr diet on control mice, and the effect of *Pparg*^ΔHep^ on HFC+Fr-fed mice, and two-way analysis of variance (ANOVA) with a post-hoc Bonferroni multiple comparisons test to assess the effect of HFC+Fr diet and *Pparg*^ΔHep^ on body composition. Metabolic cage data from cohort #4 was analyzed with CalR2 software (Metabolic Core, Beth Israel Deaconess Medical Center, Boston, MA) on the basis of total mass. To assess the effect of HFC+Fr diet or *Pparg*^ΔHep^ on overall averages of light and dark cycles (total of 96 hours of measurement) we used Student’s *t test*. To determine whether the effects on metabolic parameters across the light and dark cycles were independent of mass, we used ANCOVA. *p-*values < 0.05 were considered significant.

## Results

### HFC+Fr diet promotes steatohepatitis and maintains metabolic dysfunction promoted by the HF diet

We previously reported that feeding a HFCF diet for 16 weeks reduced adiposity and insulin resistance in mice with pre-existing HF diet-induced obesity (11). This effect may have been attributed to a modest amount of fat-derived caloric content of this HFCF vs HF diet (4.5 cal/g vs 5.2 cal/g) and the presence of trans-fats from the partially hydrogenated corn oil, which could reduce adiposity and insulin resistance in obese mice that are fed a HF diet with 60% Kcal from fat (23). In this study, we compared the early effects that feeding a HFCF diet or a HFC+Fr diet without trans-fat for 8 weeks have on body composition and development of MASH in mice with pre-existing HF diet-induced obesity. Furthermore, the effect of hepatocyte *Pparg* expression on the early onset of MASLD/MASH induced by HFCF or HFC+Fr diets has been tested with control (*Pparg*-intact mice), and adult-onset *Pparg*^ΔHep^ mice (Cohorts #1 and #2).

Eighteen weeks of a HF diet increased body weight in male mice (Fig 1A), and an extension of eight weeks of HF diet promoted body weight gain in control and *Pparg*^ΔHep^ mice (Fig 1B). Of note, eight weeks of HFCF diet prevented body weight gain in control and *Pparg*^ΔHep^ mice due to the negative effect of HFCF diet on whole-body fat mass and fat-free fluid mass (Fig. 1B-E). Similar to the HF diet, eight weeks of HFC+Fr diet promoted body weight gain increasing fat mass (Fig 1. F-J). The negative effect of the HFCF diet on adiposity of mice with HF-induced obesity seems to be selective of the amount of subcutaneous white adipose tissue and brown adipose tissue, since the levels of urogenital or mesenteric white adipose tissue were not significantly reduced by the HFCF diet. This negative effect of the HFCF diet on adipose tissue weight was not observed in mice that were switched to a HFC+Fr diet, suggesting that this HFC+Fr diet maintained obesity and metabolic dysfunction in mice with pre-existing diet-induced obesity (Fig. 1K-N). However, both diets had similar effects on plasma lipid and insulin levels, and on blood glucose level (Fig 1 O-S). While the HF diet consistently increased plasma cholesterol and insulin levels, and reduced plasma NEFA levels, HFCF and HFC+Fr diets increased plasma cholesterol and reduced plasma insulin levels in control mice with HF-induced obesity (Fig. 1. P,R). Finally, *Pparg*^ΔHep^ did not have a significant effect on the body composition changes induced by HFCF or HFC+Fr diets. However, it prevented the increase of plasma cholesterol levels and increased blood glucose in HFC+Fr-fed mice (Fig. 1P,S).

**Figure 1.**
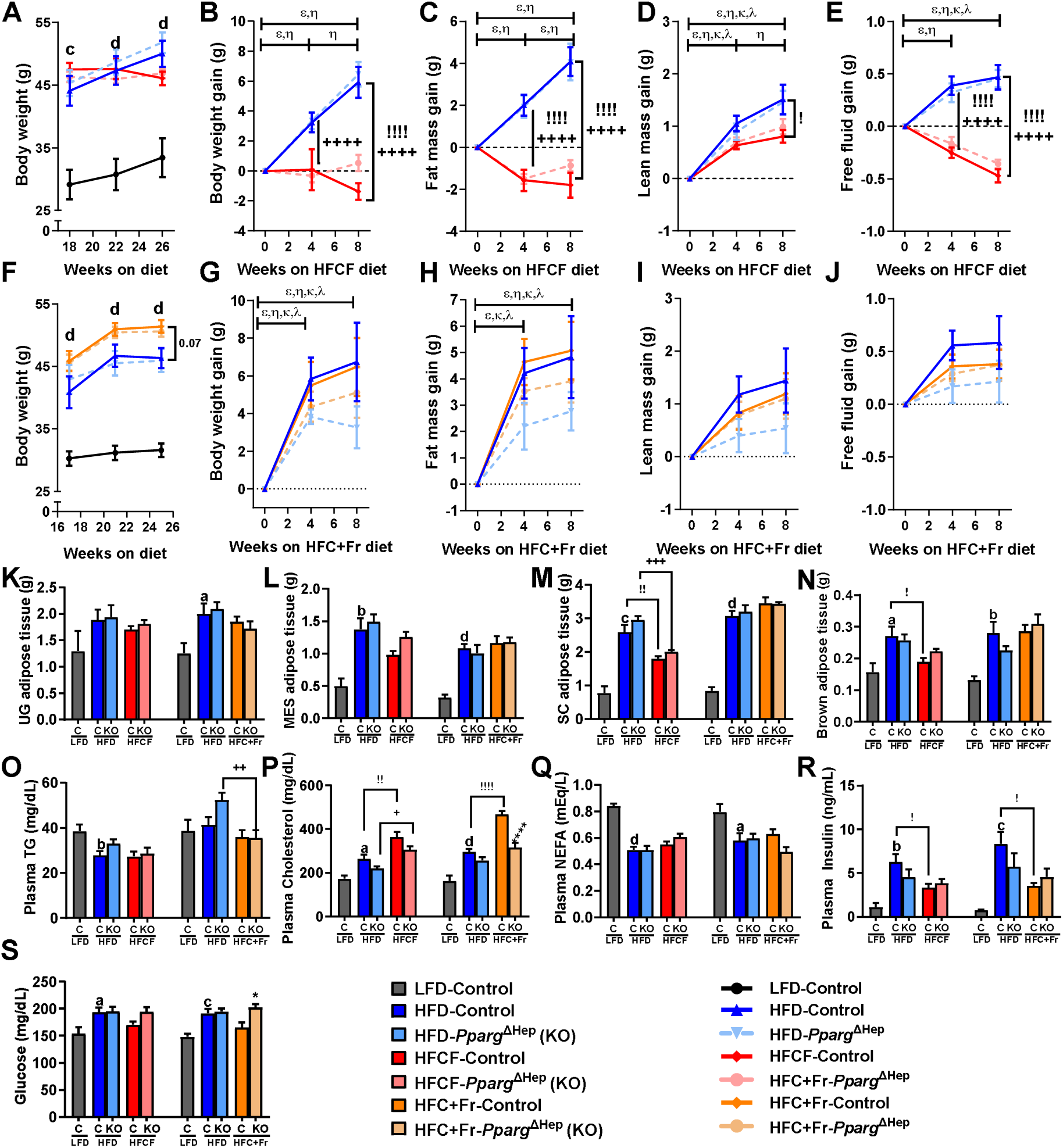
Effect of eight weeks of HFCF diet or HFC+Fr diet on body composition and metabolic endpoints of mice with pre-existing HF diet-induced obesity. Body weight progression between 18 weeks and 26 weeks of special diets in cohort #1 (A), and changes in body weight (B), fat mass (C), lean mass (D), and free fluid mass (E) during the 8 weeks of HFCF diet from mice of cohort #1. Body weight progression between 18 weeks and 25 weeks of special diets in Cohort #2 (F), and changes in body weight (G), fat mass (H), lean mass (I), and free fluid mass (J) during the 8 weeks of HFC+Fr diet from mice of cohort #2. Weight of urogenital (K, UG), mesenteric (L, MES), subcutaneous (M, SC), and brown (N, BAT) adipose tissue from mice in cohorts #1 and #2. Plasma levels of TG (O), cholesterol (P), NEFA (Q), and insulin (R), and blood glucose levels (S) from mice in cohort #1 and #2. Data are shown as mean ± standard error of the mean (SEM). Letters (a-d) indicates significant differences between LF-fed and HF-fed control mice. Greek symbols indicate significant differences (p<0.05) between 0-4 and 4-8 or 0-8 weeks between HF-control mice (ε), HF-*Pparg*^ΔHep^ mice (η), HFCF-control mice (κ), HFCF-*Pparg*^ΔHep^ mice (λ), HFC+Fr-control mice (κ), HFC+Fr-*Pparg*^ΔHep^ mice (λ). Exclamation marks (!) indicate significant differences between HF-fed and HFCF-control mice in Cohort #1 or HF-fed and HFC+Fr-control mice in Cohort #2. Plus signs (+) indicate significant differences between HF-fed and HFCF-*Pparg*^ΔHep^ mice in Cohort #1 or HF-fed and HFC+Fr-*Pparg*^ΔHep^ mice in Cohort #2. Asterisks indicate significant differences between control and *Pparg*^ΔHep^ mice. a, !, * p<0.05; b, !!, ++, p<0.01; c, +++, p<0.001; d, !!!!, ++++, ****, p<0.0001. n= 5-11 mice/group in cohort #1 and 7-8 mice/group in cohort #2.

Next, we compared the impact that feeding an HFCF or HFC+Fr diet for eight weeks has on the liver of mice with pre-existing diet-induced obesity. HF diet-induced obesity was associated with increased liver weight and steatosis (liver TG) as compared to control mice fed a LF diet, and eight weeks of HFCF diet further increased liver weight and TG as compared to their HF-fed counterparts (Fig. 2A-B). Moreover, the HFCF diet increased liver cholesterol content and plasma ALT levels that were not increased by the HF diet (Fig. 2C-D). Similarly, the HFC+Fr diet increased liver weight, cholesterol, and plasma ALT, without a significant increase in liver TG content (Fig. 2A-D). These data indicate that HFCF and HFC+Fr diets contribute to the progression of MASLD. In fact, the histological analysis of H&E-stained liver sections showed that the HF diet increased steatosis (Supplementary Figure 1), and both HFCF and HFC+Fr diets did not increase steatosis further than the HF diet did, but increased hepatocyte ballooning and inflammation (Supplementary Figure 1) to promote MASH (MASLD activity score >5, Fig. 2E, G,H). Interestingly, only 8 weeks of HFC+Fr diet, but not HFCF diet, in mice with pre-existing diet-induced obesity promoted the development of MASH with fibrosis (Fig. 2F, H), suggesting that this new HFC+Fr dietary approach could be a new MASH-inducing diet. Of note, *Pparg*^ΔHep^ significantly reduced liver weight in HFCF- and HFC+Fr-fed mice, and liver TG in the HF-fed mice of the cohort #1. Interestingly, *Pparg*^ΔHep^ mice had a significant effect on reducing plasma ALT levels, MASLD activity score, and fibrosis in HFC+Fr-fed mice (Fig. 2D, E, F). The histological effects of these diets were associated with alteration of hepatic gene expression. The activity of hepatocyte PPARγ must be increased in HFCF and HFC+Fr-fed mice as suggested by the increased expression of the PPARγ-target gene *Cidec*. However, the knockout of *Pparg* gene in *Pparg*^ΔHep^ mice dramatically reduced the expression of hepatic *Cidec* (Fig. 2I,J). To assess the development of MASH by eight weeks of HFCF and HFC+Fr diet in mice with pre-existing diet-induced obesity we confirmed the increased expression of *Col1a1, Timp1, Tnfa, Ccl2,* and *Trem2* in these mice (Fig. 2K-O) as we have reported in male mice fed a HFCF diet for 24 weeks (16) In our previous studies, we reported that HFCF diet reduces the expression of genes involved in methionine metabolism (11,16). Only 8 weeks of HFCF and HFC+Fr diet in mice with pre-existing diet-induced obesity reduced the expression of hepatic *Gnmt* and *Pemt*, and HFC+Fr diet reduced the expression of *Bhmt (*Fig. 2P-R). Overall, the effects of HFCF and HFC+Fr diets on the regulation of these hepatic genes were similar to those published by our group previously. In addition, *Pparg*^ΔHep^ reduced the expression of *Tnfa* and *Ccl2* in HFCF-fed mice, and that of *Timp1* and *Ccl2* in HFC+Fr-fed mice, whereas increased the expression of *Gnmt* and *Bhmt* in HFC+Fr-fed mice. These effects of *Pparg*^ΔHep^ may indicate that in the absence of hepatocyte *Pparg* expression, the development of diet-induced MASH is slowed down.

**Figure 2.**
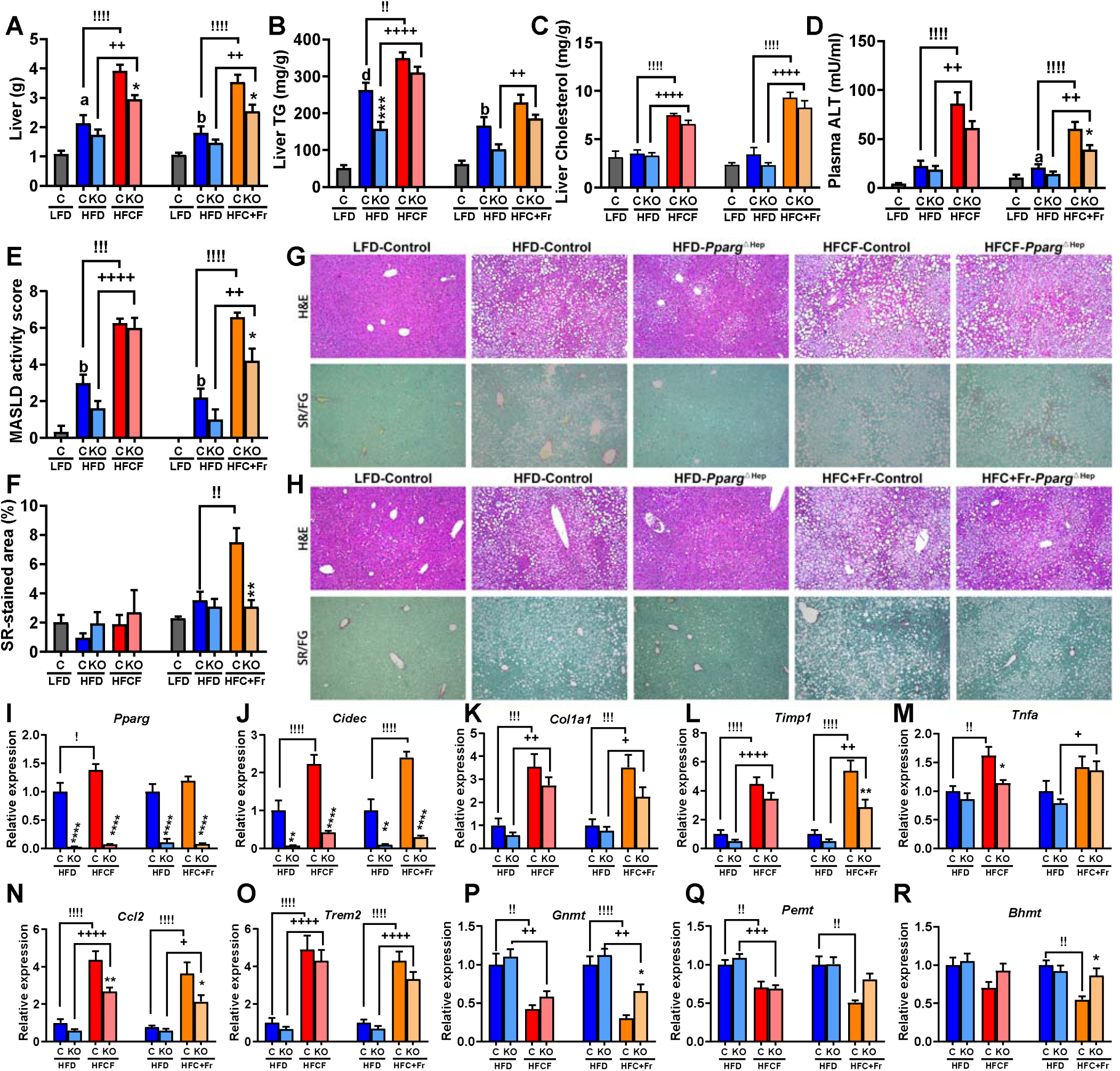
MASH-inducing effects of eight weeks of HFCF diet or HFC+Fr diet on mice with pre-existing HF diet-induced obesity. Liver weight (A), liver TG content (B), liver cholesterol content (C), plasma ALT level (D), MASLD activity score (E), and picrosirius red-stained fibrotic area (F) of mice in cohorts #1 [LF, HF, and HFCF-fed mice] and #2 [LF, HF, and HFC+Fr-fed mice]. Representative pictures of hematoxylin & eosin (H&E)- and picrosirius red/fast green (SR/FG)-stained liver section of mice in cohort #1 (G, LF, HF, and HFCF-fed mice) and cohort #2 (H, LF, HF, and HFC+Fr-fed mice). Hepatic gene expression of *Pparg* (I), *Cidec* (J), *Col1a1* (K), *Timp1* (L), *Tnfa* (M), *Ccl2* (N), *Trem2* (O), *Gnmt* (P), *Pemt* (Q), and *Bhmt* (R) of mice in cohort #1 (HF, and HFCF-fed mice) and cohort #2 (HF, and HFC+Fr-fed mice). I-R, expression level is represented as relative values of HF-fed control mice in cohort #1 or coho rt #2. Data are shown as mean ± standard error of the mean (SEM). Letters (a-d) indicate significant differences between LF-fed and HF-fed control mice. Exclamation marks (!) indicate significant differences between HF-fed and HFCF-control mice in Cohort #1 or HF-fed and HFC+Fr-control mice in Cohort #2. Plus signs (+) indicate significant differences between HF-fed and HFCF-*Pparg*^ΔHep^ mice in Cohort #1 or HF-fed and HFC+Fr-*Pparg*^ΔHep^ mice in Cohort #2. Asterisks indicate significant differences between control and *Pparg*^ΔHep^ mice. a, !, * p<0.05; b, !!, ++, ** p<0.01; !!!,+++, *** p<0.001; d, !!!!, ++++, ****, p<0.0001. n= 5-11 mice/group in cohort #1 and 7-8 mice/group in cohort #2.

### Obesogenic and sex-dependent hepatic effects of HFC+Fr diet in male and female mice

We fed male and female control and *Pparg*^ΔHep^ mice for 24 weeks an HFC+Fr diet to assess whether HFC+Fr diet induces obesity, insulin resistance, and MASH with fibrosis. In male mice, HFC+Fr diet induced a rapid increase in body weight that reached 56.58 +/- 1.27g after 24 weeks of diet, which was mostly due to an increase in fat mass rather than lean mass and free fluid mass (Fig. 3A-E). Female mice fed an HFC+Fr diet for 24 weeks weighed 46.31g +/- 2.33g, and their body weight gain was also mostly due to an increase in fat mass (Fig. 3F-J). However, the body weight and fat mass gain in female mice were slower than that observed in male mice, and continued to increase by 24 weeks of diet. In both male and female mice, the increased fat mass was associated with an increase in white and brown adipose tissue, with the exception of urogenital fat mass in male mice (Fig. 3 K-O). As described above in the cohort #2, the HFC+Fr diet did not increase plasma TG levels as compared to mice fed a LF diet, but it increased plasma cholesterol levels and reduced plasma NEFA levels in both male and female mice (Fig. 3 P-R). Furthermore, the HFC+Fr diet significantly increased plasma insulin levels in male and female mice, as well as blood glucose in female mice, which was associated with glucose intolerance and increased fasting blood glucose after 21 weeks of the HFC+Fr diet in both male and female mice (Fig. 3.S-W). In another cohort of male mice, we assessed metabolic rate. After 16 weeks of diet, body weight of HFC+Fr-fed mice was 44.83 +/- 1.57g, and HFC+Fr-fed mice showed reduced the respiratory exchange ratio in the dark and light cycles which was indicative of a preferential use of lipids as source of energy (Fig. 4A,B). Average values of energy expenditure, food consumption, oxygen consumption, and carbon dioxide production showed clear diurnal variations in LF- and HFC+Fr-fed mice. Of note, HFC+Fr diet increased energy expenditure and oxygen consumption (Fig 4 C-F, Supplemental Figure 2). However, the ANCOVA analysis indicated that, independent of mass, energy expenditure and oxygen consumption were significantly increased by HFC+Fr only during the light cycle (Fig 4C-F, Supplemental Figure 3). Interestingly, *Pparg*^ΔHep^ had a minimal impact on the obesogenic effects of HFC+Fr diet, and only reduced plasma cholesterol in male mice, increased plasma NEFA in female mice, and plasma insulin in male mice (Fig. 3Q-S). Interestingly, *Pparg*^ΔHep^ also increased energy expenditure and oxygen consumption in HFC+Fr-fed male mice during the light cycle (Fig 4C-F, Supplemental Figure 3) but without altering obesity or the assessed metabolic endpoints.

**Figure 3.**
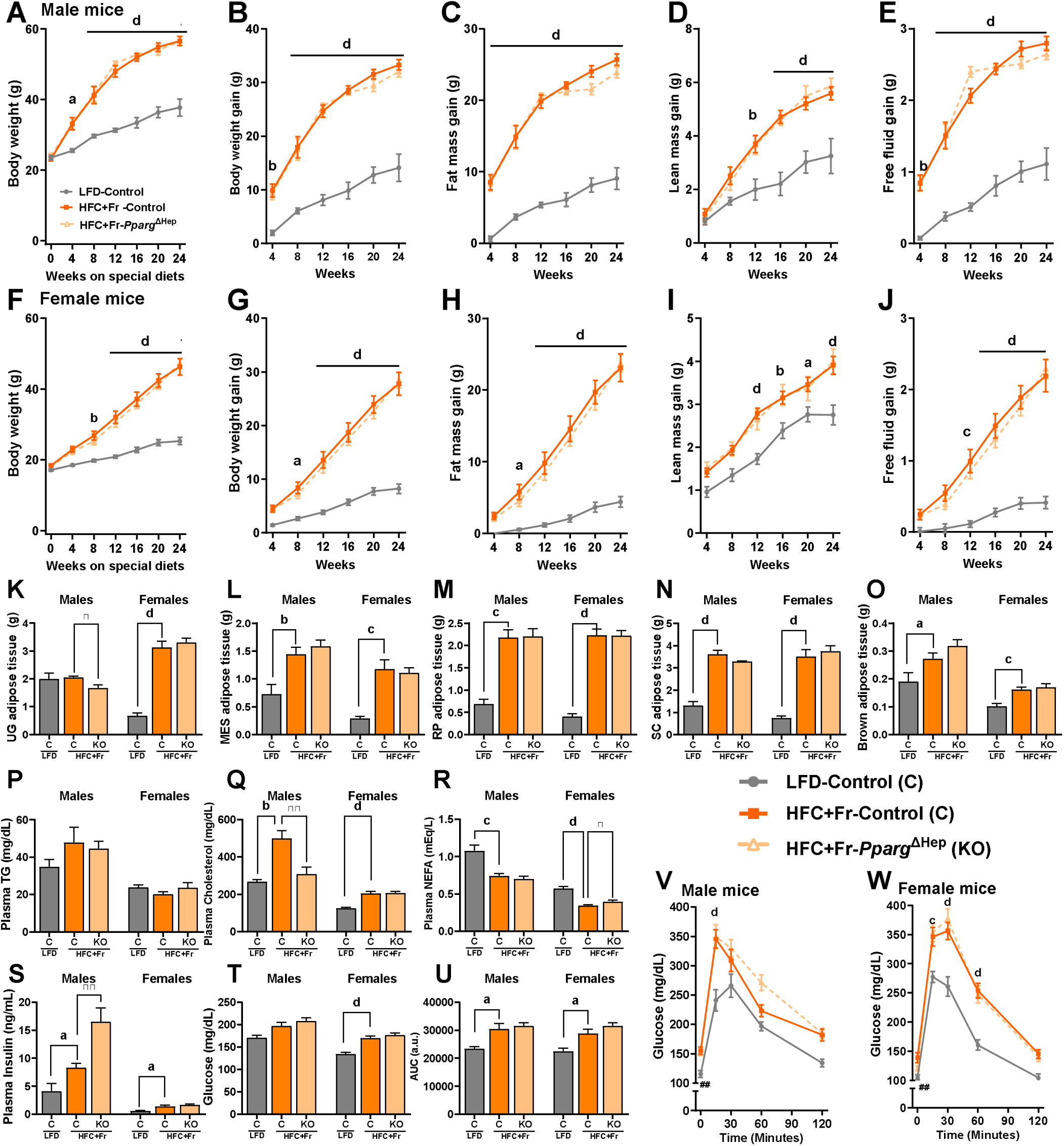
Effect of twenty-four weeks of HFC+Fr diet on body composition and metabolic endpoints of male and female mice. Body weight progression (A), and changes in body weight (B, BW), fat mass (C), lean mass (D), and free fluid mass (E) during the 24 weeks of HFC+Fr diet in male mice. Body weight progression (F), and changes in body weight (G, BW), fat mass (H), lean mass (I), and free fluid mass (J) during the 24 weeks of HFC+Fr diet in female mice. Weight of urogenital fat (K, UG-fat), mesenteric fat (L, MES-fat), retroperitoneal fat (M, RP-fat). subcutaneous fat (N, SC-fat), brown adipose tissue (O, BAT) from male and female mice. Plasma levels of TG (P), cholesterol (Q), NEFA (R), insulin (S), and blood glucose levels (T) from male and female mice. Area under the curve (U, AUC), and glucose tolerance test of male (V) and female (W) mice. Data are shown as mean ± standard error of the mean (SEM). Letters (a-d) indicate significant differences between LF-fed and HFC+Fr-fed control mice. Asterisks indicate significant differences between HFC+Fr-fed control and HFC+Fr-fed *Pparg*^ΔHep^ mice. # indicate significant differences between LF-fed control and HFC+Fr-fed control in t=0 of the glucose tolerance test. a, * p<0.05; b, **, ## p<0.01; c, p<0.001; d, ****, p<0.0001. n= 4-9 male mice/group and 8-11 female mice/group.

**Figure 4.**
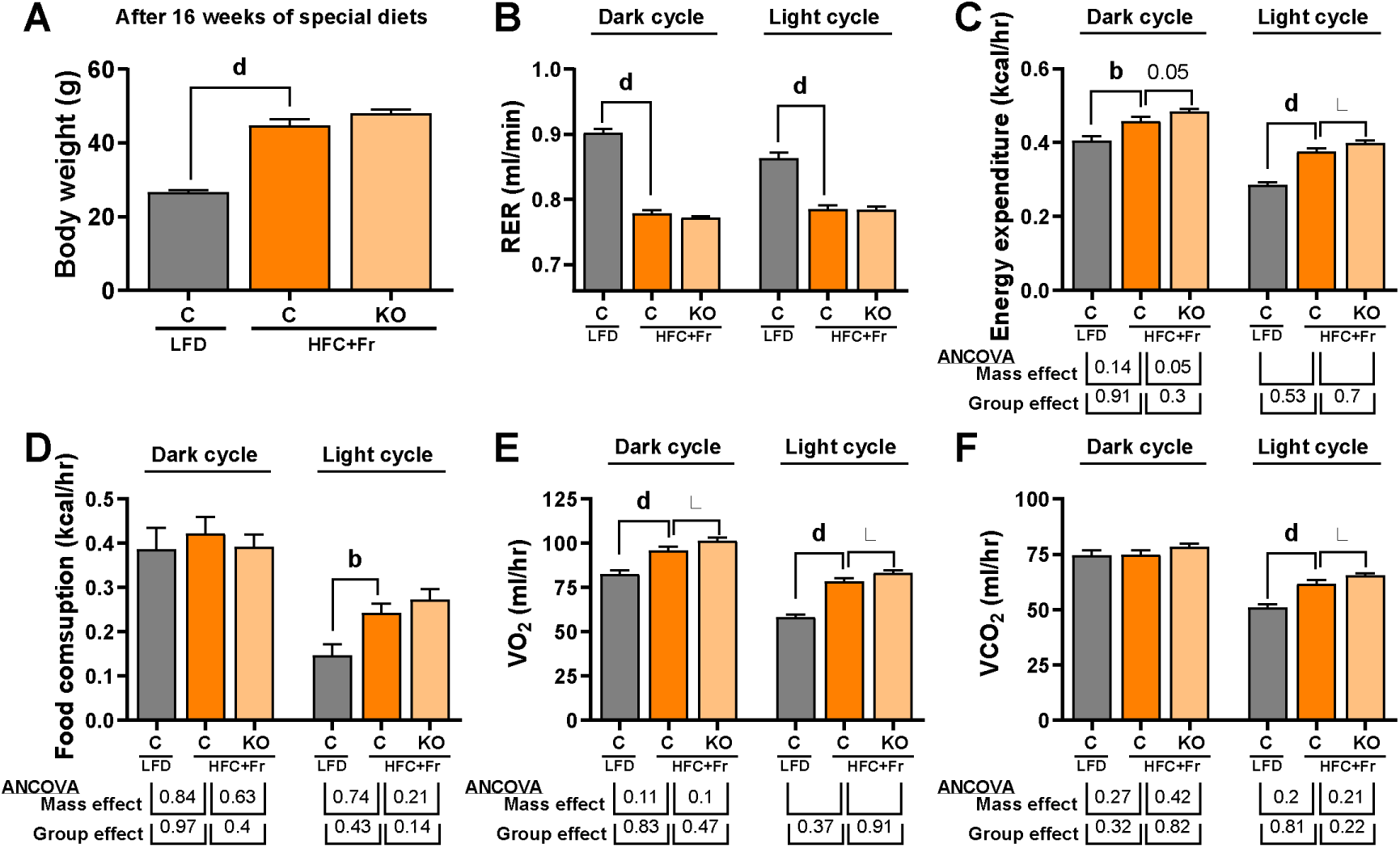
Effect of sixteen weeks of HFC+Fr diet on the metabolic rate of male mice. Body weight (A), respiratory exchange ratio (B, RER), energy expenditure (C), food consumption (D), volume of oxygen consumption (E), and volume of carbon dioxide production (F). Data are shown as mean ± standard error of the mean (SEM). Letters (b-d) indicate significant differences between LF-fed and HFC+Fr-fed control mice. Asterisks indicate significant differences between HFC+Fr-fed control and HFC+Fr-fed *Pparg*^ΔHep^ mice. * p<0.05; b, p<0.01; d, p<0.0001. n= 11-13 mice/group

Next, we compared the effects of 24 weeks of HFC+Fr diet on the liver of male and female mice. HFC+Fr diet increased liver weight, steatosis (liver TG and cholesterol), plasma ALT levels, and MASLD activity score in both male and female mice (Fig. 5A-E). Of note, the histological analysis of H&E-stained liver sections showed that the HFC+Fr diet dramatically increased steatosis, hepatocyte ballooning, and inflammation in male mice and to a lesser extent in female mice (Fig. 5H,I, and Supplementary Figure 4). Notably, the average MASLD activity score was greater than 5 (MASH) in male mice, which was associated with increased fibrosis, as indicated by the picrosirius red-stained area of liver sections and collagen 1A1 levels (Fig. 5E-I). We have previously published that hepatic *Pparg* expression is increased in mice with MASH and fibrosis, as well as in patients with obesity and MASLD/MASH (11,16). Of note, HFC+Fr diet increased the expression of hepatic *Pparg* gene and PPARγ2 in both male and female mice (Fig 5J-K). *Pparg*^ΔHep^ significantly reduced the expression of *Pparg* gene and that of PPARγ1 and PPARγ2 in both male and female mice (Fig 5K). This was associated with reduced liver steatosis in HFC+Fr-fed male and female mice (Fig. 5B), and reduced liver weight, plasma ALT, MASLD activity score, and fibrosis quantified by picrosirius red-stained liver section area and Collagen 1 protein level in HFC+Fr-fed male mice (Fig. 5A,D-G). To explore if hepatic *Ppar*γ expression is associated with the onset of steatosis, inflammation, and fibrosis in male and female mice, we assessed the expression of *Pparg*-target genes *Cidea*, *Cidec*, pro-inflammatory genes: *Ccl2* and *Trem2*, and pro-fibrogenic genes: *Col1a1* and *Timp1*, in the livers of HFC+Fr-fed mice. HFC+Fr diet increased the expression of these genes in males, but not females. Of note, *Pparg*^ΔHep^ reduced the expression of *Pparg*-traget genes: *Cidea* and *Cidec* in male mice. It also reduced the expression of *Col1a1* and *Timp1* in male mice, as previously reported in mice that were fed a HFCF diet for 24 weeks (16), (Fig. 5L-Q). In addition, the HFC+Fr diet reduced the expression of genes involved in hepatic methionine metabolism: *Mat1a*, *Gnmt*, *Pemt, Ahcy*, and *Bhmt* but not *Cbs* in male mice, and *Pparg*^ΔHep^ prevented the negative effect of HFC+Fr diet on *Gnmt, Pemt,* and *Bhmt* (Fig. 5R-W). We further confirmed that *Pparg* is activated in HFC+Fr-fed mice, and is negatively associated with methionine metabolism, as indicated by the protein levels of hepatic CIDEC (a PPARγ-target gene) and BHMT (Fig. 5X,Y). Again, the data from this third cohort of mice confirmed the association between hepatocyte *Pparg* expression and features of diet-induced MASLD/MASH, as shown above male mice (Figure 2) and in our previous reports using male mice (11,16). Furthermore, these data indicate that the obesogenic effects of HFC+Fr diet also has a *Pparg*-dependent negative effects on hepatic methionine metabolism in male mice that could promote MASH and fibrosis.

**Figure 5.**
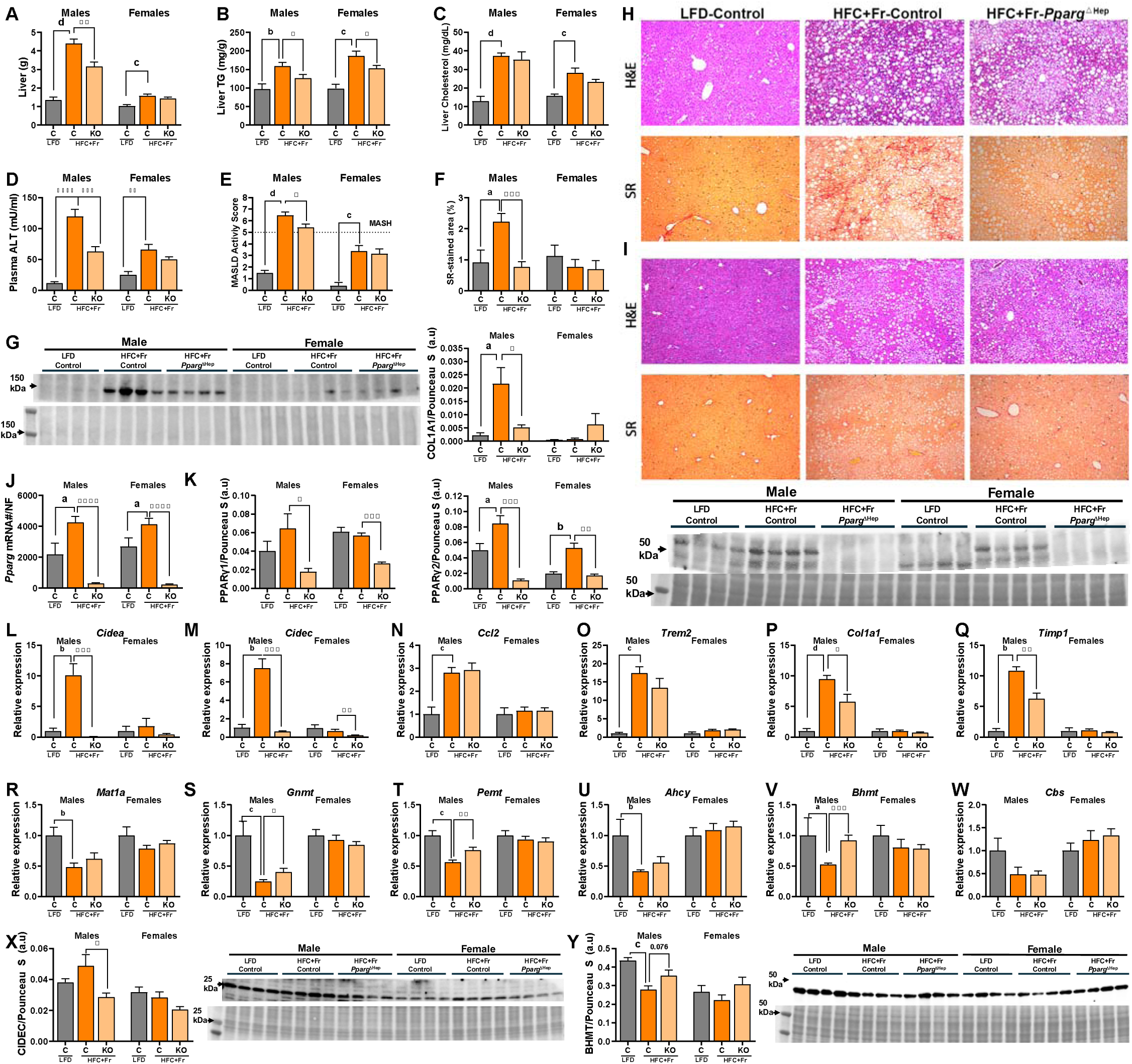
MASH-inducing effects of twenty-four weeks of HFC+Fr diet on male and female mice. Liver weight (A), liver TG content (B), liver cholesterol content (C), plasma ALT level (D), MASLD activity score (E), picrosirius red-stained fibrotic area of liver sections (F, SR), western blot for hepatic COL1A1 (G, top) with its Ponceau S staining (G, bottom) and quantification from male and female mice (G). Representative pictures of hematoxylin & eosin (H&E)- and picrosirius red (SR)-stained liver section of male (H) and female (I) mice. Hepatic gene expression of *Pparg* mRNA (J) and western blot for hepatic PPARγ (K, top) with its Ponceau S blot (K, bottom) and quantification of PPARγ1 and PPARγ2 from male and female mice (K). Hepatic gene expression of *Cidea* (L), *Cidec* (M), *Ccl2* (N), *Trem2* (O), *Col1a1* (P), *Timp1* (Q), *Mat1a* (R), *Gnmt* (S), *Pemt* (T), *Ahcy* (U), *Bhmt* (V), *Cbs* (W) of male and female. L-W, the expression level is represented as relative values of LF-fed control mice. Western blots for hepatic CIDEC (X, top) and BHMT (Y, top) with their Ponceau S staining (X, Y, bottom) and quantification from male and female mice. Data are shown as mean ± standard error of the mean (SEM). Letters (a-d) indicate significant differences between LF-fed and HFC+Fr-fed control mice. Asterisks indicate significant differences between HFC+Fr-fed control and HFC+Fr-fed *Pparg*^ΔHep^ mice. a, * p<0.05; b, **, p<0.01; c,*** p<0.001; d, ****, p<0.0001. n= 4-9 male mice/group and 8-11 female mice/group.

To further characterize the hepatic phenotype induced by the HFC+Fr diet, we performed RNAseq on liver samples from male and female mice. The analysis of the RNAseq confirmed the strong effect of the HFC+Fr diet on male mice with 2049 differential expressed genes (DEG, Log2FC <-1, >1, padj <0.05), as compared to the 157 DEG in female mice (Fig. 6A). Although hepatic *Pparg* was increased in both HFC+Fr-fed male and female mice, *Pparg*^ΔHep^ differentially regulated the expression of 238 genes in HFC+Fr-fed male mice and only 12 in HFC+Fr-fed female mice (Fig. 6B). A heatmap of HFC+Fr-regulated PPARγ-target genes and genes involved in hepatic inflammation, fibrosis and methionine metabolism confirmed the results obtained by qPCR and support the sex-specific effect of HFC+Fr diet on PPARγ activation and regulation of genes associated with onset of MASH (Fig 6. C,D). Overall, these findings support the sex-dependent effect of the HFC+Fr diet in the regulation of hepatic transcriptomics, and confirm that hepatocyte PPARγ is a relevant regulator of hepatic transcriptomics in HFC+Fr-fed male mice. The enrichment analysis of the transcriptomic data showed that the HFC+Fr diet upregulated gene ontology (GO) terms related with inflammation and immune response in male mice, whereas it downregulated those involved in methionine metabolism and mitochondrial function, as we reported in previous studies with the HFCF diet (Fig. 6E, left). In female mice, the HFC+Fr diet upregulated GO terms related to amino acid metabolism and reduced some GO terms related with immune response and lipid metabolism (Fig. 6E, right). *Pparg*^ΔHep^ upregulated GO terms and pathways associated with amino acid metabolism and downregulated of those involved in immune response in male mice (Fig. 6F, left), and upregulated GO terms related to immune defense and downregulated those involved in angiogenesis and lipid metabolism in female mice (Fig 6F, right).

**Figure 6.**
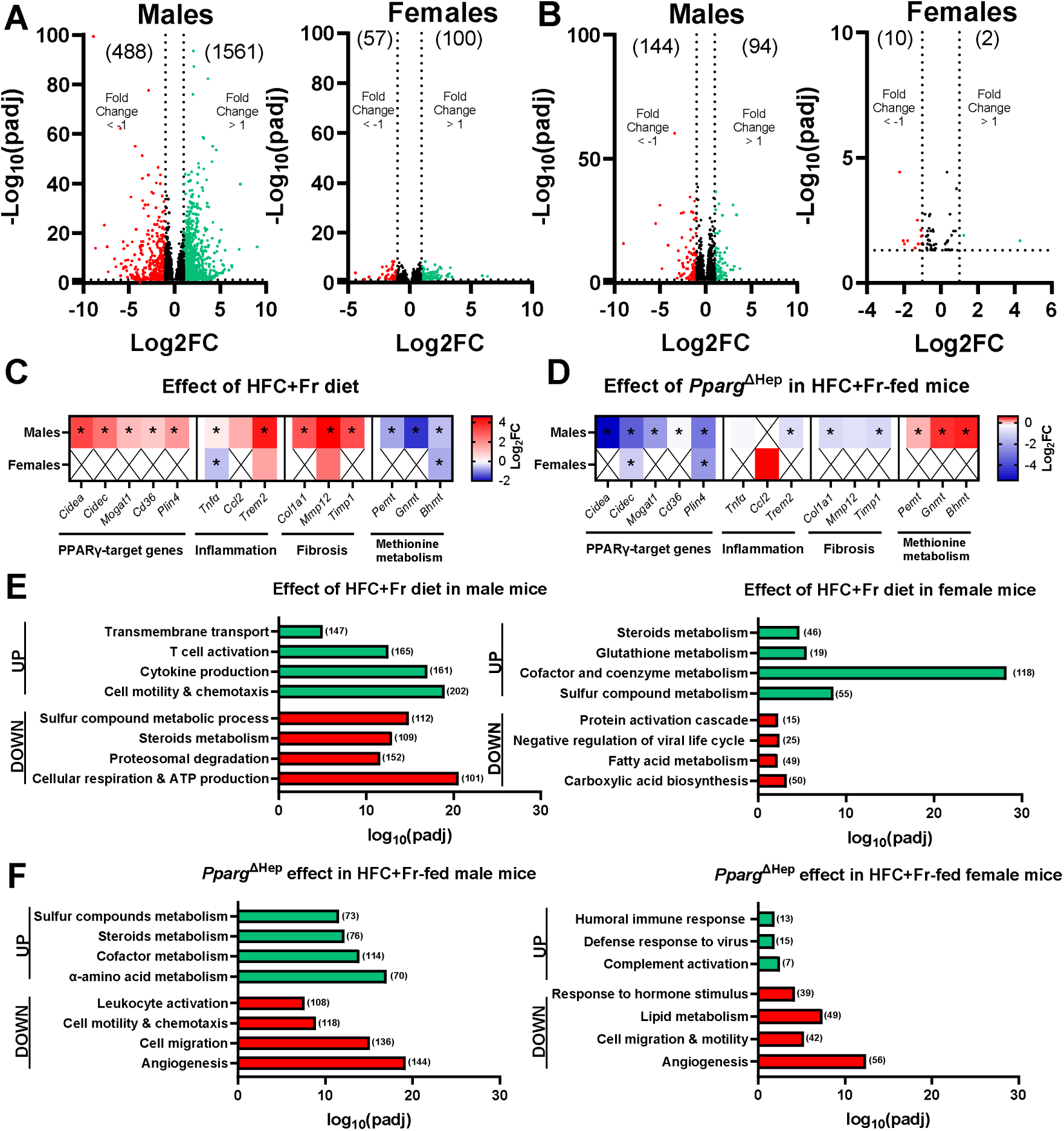
Effect of twenty-four weeks of HFC+Fr diet and *Pparg*^ΔHep^ on hepatic transcriptome of mice. Volcano plots showing the differentially expressed genes (DEG, padj<0.05) by HFC+Fr diet (A) and *Pparg*^ΔHep^ (B) in male and female mice. Heatmaps showing the regulation of PPARγ-target genes: *Cidea*, *Cidec*, monoacylglycerol O-acyltransferase 1 (*Mogat1*), fatty acid translocase (*Cd36*), perilipin 4 (*Plin4*), the inflammation-related genes: *Tnfa*, *Ccl2*, *Trem2*, the fibrosis-related genes: *Col1a1*, metalloproteinase 12 (*Mmp12*), *Timp1*, and the methionine metabolism-related genes: *Pemt*, *Gnmt*, *Bhmt*, by HFC+Fr diet (C) and *Pparg*^ΔHep^ (D) in male and female mice. Asterisk (p<0.05) indicate significant regulation. Selected groups of genes identified by the enrichment analysis of DEG by HFC+Fr diet (E) and *Pparg*^ΔHep^ (F) in male and female mice. The number of upregulated (green) or downregulated (red) DEG in the volcano plots or enrichment analysis are indicated in brackets. n= 4-6 male mice/group and 5 female mice/group.

Taken together, HFC+Fr is an obesogenic diet that increases liver weight and steatosis in both male and female mice, which could be used as a more reliable tool for studying obesity-driven MASLD/MASH in mice. However, the HFC+Fr diet promotes MASH and fibrosis only in male mice, and hepatocyte *Pparg* contributes to the progression of liver disease by regulating methionine metabolism and fibrosis in the liver. In contrast, female mice did not develop liver disease under the same dietary conditions, despite showing increased hepatocyte *Pparg* expression.

## Discussion

A major challenge in studying the molecular mechanisms driving the progression of human-like MASLD to MASH is the generation of an experimental model that recapitulates steatohepatitis with fibrosis and peripheral metabolic dysfunction with a dietary approach in young adult/middle aged mice (6-10 months). Several decades ago, special diets deficient in choline or methionine were used to rapidly induce fatty liver with fibrosis (24). A lack of choline supply reduces TG export from hepatocytes as VLDL, and reduced methionine supply induces cellular stress, driving hepatic inflammation and fibrosis (25,26). However, choline and methionine restriction reduce adiposity and circulating lipid levels, and improve insulin sensitivity (7). Therefore, these nutrient-deficient diets cannot reproduce human-like MASLD/MASH with obesity and insulin resistance in animal models. By contrast, HF diets have been extensively used to induce steatosis, obesity, dyslipidemia, and insulin resistance (27). However, most HF diets fail to induce the progression from steatosis to MASH with fibrosis (11,28) unless that they are provided chronically (>50 weeks (29)), which also limits their use to study advanced stages of MASLD and MASH in young adult/middle aged mice. To overcome this limitation, HF diets containing cholesterol, fructose, and various sources of fat, including those containing trans fats, were developed. Feeding these diets for six months induces steatohepatitis with marked fibrosis and metabolic dysfunction in male mice (28,30). However, when these diets are provided alongside nutrient-matched low-fat control diets, it becomes evident that they do not induce severe adiposity as HF diets do (23,31). Previously, we used a new version of HF diet with cholesterol and fructose that contains partially hydrogenated corn-oil as source of trans fat (a version of AMLN-diet). This HFCF diet induces MASH with fibrosis with a modest effect on body weight gain in male mice (16), and when it is provided to mice with pre-existing diet-induced obesity, it promotes MASH with fibrosis but reducing adiposity and insulin resistance (11). This negative effect of the HFCF diet on metabolic dysfunction limits the translational potential of the HFCF-fed mice as a model of human-like MASH. In this study, we reproduced the early negative effect of an HFCF diet on adiposity in mice with pre-existing diet-induced obesity (11) and showed that only 8 weeks of HFCF diet increased liver weight and promoted MASH progression in these mice with pre-existing HF diet-induced obesity. Interestingly, a new formulation of HF diet with 60% Kcal from fat (mainly lard) and 2% cholesterol, but supplemented with fructose in the drinking water (HFC+Fr diet) did not reduce adiposity in mice with pre-existing obesity, but induced MASH with significant fibrosis in male mice. Of note, this HFC+Fr diet does not contain trans-fat as the fat found in the HFCF diet, nor palm oil, which is used in the Gubra-Amylin NASH (GAN) diet (10). Thus, the use of lard as a fat source in the HF diet, plus the addition of cholesterol in the diet and fructose in the drinking water, is a key component in inducing MASH and sustaining metabolic dysfunction. Overall, this study confirms that 24 weeks of an HFC+Fr diet efficiently induces obesity and metabolic dysfunction associated with steatosis in both male and female mice, and this MASLD phenotype progresses to MASH with fibrosis in male mice.

HFC+Fr diet induced MASLD with obesity in both sexes; however the MASH-inducing effects of HFC+Fr diet were sex- and hepatocyte PPARγ-dependent. In fact, hepatocyte *Pparg* expression associated with the development of HF diet-induced steatosis (13,14,32,33) contributed to the progression of HFCF diet-induced MASH with fibrosis only in male mice (11,16). Although hepatocyte *Pparg* mRNA and hepatic PPARγ protein was increased in both HFC+Fr-fed male and female mice, the transcriptional activity of PPARγ may be sex-dependent because the PPARγ-target genes *Cidea* and *Cidec* were only increased in the livers of male mice. This was in line with our previous report, in which hepatic *Cidea* and *Cidec* expression were increased in HF-fed male but not female control (*Pparg*-intact) mice (11). Also, it should be noted that *Cidea* and *Cidec* are negatively regulated by ovarian hormones in a *Ppargc1a*-dependent manner (34,35). In fact, estrogen signaling interferes with PPARγ transcriptional activity in adipose tissue and breast cancer cells (36,37), so in females, estrogen signaling in hepatocytes may reduce PPARγ transcriptional activity. Furthermore, we confirmed that the knockout of hepatocyte PPARγ reduced the expression of CIDEC protein in the liver of male mice in this study. PPARγ-regulated CIDEC (38) promotes the storage of TG (39,40) and the development of steatohepatitis in mice and humans (41), which could explain the association of PPARγ and CIDEC with the progression of MASH in HFC+Fr-fed male mice. Interestingly, the hepatic transcriptome of mice that were fed a HFC+Fr diet for 24 weeks was also dramatically altered in male mice but not in females. We have previously shown that 16 weeks of HFCF diet dramatically altered the hepatic transcriptome of both male and female mice with pre-existing HF diet-induced obesity, and increased the expression of PPARγ-target genes, and genes involved in inflammation and fibrosis. However, in that study, both male and female mice progressed to MASH with a significant upregulation of inflammatory and fibrogenic genes (11). In the current study, *Pparg*^ΔHep^ reduced steatosis in both HFC+Fr-fed male and female mice, but only reduced MASLD activity score (MASH), fibrosis, plasma ALT levels, and PPARγ-target genes in male mice. This sex-dependent effect on the regulation of key PPARγ-regulated genes in the liver of male mice could be associated with the modest regulation of hepatic transcriptome in female mice fed an HFC+Fr diet. Nevertheless, it is possible that if HFC+Fr diet would have been provided for more than 24 weeks, as we did in our previous study with the combination of HF and HFCF diets, that had negatively impacted ovarian function and promoted the development of diet-induced MASH (11,31,42).

We previously reported that increased hepatocyte *Pparg* expression in mice with MASH was negatively associated with hepatic regulation of methionine metabolism (11,16). In this study, the HFC+Fr diet-induced MASLD was also associated with a negative sex-dependent regulation of genes involved in hepatic methionine metabolism. The negative effect of MASLD on the expression of hepatic *Gnmt*, *Pemt*, and *Bhmt* was hepatocyte PPARγ-dependent. The downregulation of hepatic *Gnmt, Pemt,* or *Bhmt* by HFC+Fr diet in male mice is significant because the knockout of these genes induces the development of steatosis and MASLD in male mice (43–45). In our study, the upregulated hepatic expression of these genes indicates that the capacity of hepatocytes to promote S-adenosylmethionine-dependent methyltransferase activities (GNMT and PEMT) was enhanced in HFC+Fr-fed *Pparg*^ΔHep^ male mice, which could improve liver health. Furthermore, the ability to remethylate homocysteine by BHMT was not negatively impacted by HFC+Fr diet on *Pparg*^ΔHep^ male mice, and that would prevent the cellular stress induced by the increased levels of homocysteine in mice with MASLD (46,47). Therefore, hepatocyte *Pparg* may serve as a negative regulator of these genes and dysregulate methionine metabolism in the liver, thereby promoting the development of MASH (16).

Overall, this study shows that adding cholesterol to the HF diet with 60% of Kcal from fat and providing fructose in drinking water is a suitable dietary approach to induce metabolic dysfunction and steatohepatitis with fibrosis within 6 months. The obesogenic effect of this dietary approach is remarkable and stronger than that of the GAN diet, due to its caloric content and the excess carbohydrate intake provided in the drinking water, which mimics the use of fructose-containing beverages consumed alongside high-calorie food intake. Moreover, the HFC+Fr diet induced insulin resistance in both male and female mice and increased plasma cholesterol levels. However, the induction of MASH by HFC+Fr diet is sex and hepatocyte PPARγ-dependent, and the negative effects of PPARγ expression on the liver that contribute to the progression of MASH are not associated with reduced adiposity nor improved peripheral metabolic function. Our study also confirms that HFC+Fr diet-mediated progression of MASH is associated with dysregulation of methionine metabolism in the liver, and suggests that hepatocyte PPARγ may negatively impact the transmethylation of S-adenosylmethionine to methylate phospholipids and glycine, and to remethylate homocysteine which will reduce the negative effects of homocysteine accumulation in the liver (46,47). In conclusion, our study shows a new dietary approach that mimics human-like MASH associated with metabolic dysfunction and confirms the negative role of hepatocyte PPARγ in the progression of MASH in male mice, which may be associated with the dysregulation of methionine metabolism.

## Author contributions

JCC conceived and designed experiments. MSC, IH, SML, JCC, wrote the manuscript. JCC, MSC, SML, IH, JM performed experiments and analyzed data. All authors have access to the study and have reviewed and approved the final manuscript.

## Acknowledgments

Fixed samples and H&E-stained slides were processed by the Research Histology Core at the University of Illinois Chicago.

