## Supplemental Tables and Figures for "Sex- and hepatocyte PPARγ-dependent effects of an obesogenic dietary approach to induce MASH with fibrosis in mice"

**Supplemental Table 1. Composition of special diets used in this study.**

|  | LFD | | | HFD | | HFCF | | HFC+Fr | |
| --- | --- | --- | --- | --- | --- | --- | --- | --- | --- |
| **Catalog # Research Diets, Inc.** | **D12450J** | | | **D12492** | | **D16010101** | | **D12120101** | |
|  | gm% | *kcal%* | gm% | | *kcal%* | gm% | *kcal%* | *gm%* | *kcal%* |
| Protein | 19 | 20 | 26 | | 20 | 22 | 20 | 26 | 20 |
| Carbohydrate | 67 | 70 | 26 | | 20 | 45 | 40 | 26 | 20 |
| Fat | 4 | 10 | 35 | | 60 | 20 | 40 | 34 | 60 |
| Total |  | 100 |  | | 100 |  | 100 |  | 100 |
| kcal/gm | 3.8 |  | 5.2 | |  | 4.5 |  | 5.1 |  |
| **Ingredient** | **gm** | **cal** | **gm** | | **cal** | **gm** | **cal** | **gm** | **cal** |
| **Carbohydrates** |  |  |  | |  |  |  |  |  |
| Corn Starch | 506.2 | 2025 | 0 | | 0 | 0 | 0 | 0 | 0 |
| Maltodextrin 10 | 125 | 500 | 125 | | 500 | 100 | 400 | 125 | 500 |
| Sucrose | 68.8 | 275 | 68.8 | | 275 | 96 | 384 | 68.8 | 275 |
| Fructose | 0 | 0 | 0 | | 0 | 200 | 800 | 0 | 0 |
| **Fat** |  |  |  | |  |  |  |  |  |
| Soybean Oil | 25 | 225 | 25 | | 225 | 25 | 225 | 25 | 225 |
| Lard | 20 | 180 | 245 | | 2205 | 20 | 180 | 245 | 2205 |
| Corn Oil, Partially Hydrogenated | 0 | 0 | 0 | | 0 | 135 | 1215 | 0 | 0 |
| Cholesterol | 0 | 0 | 0 | | 0 | 18 | 0 | 16 | 0 |
| **Protein** |  |  |  | |  |  |  |  |  |
| Casein | 200 | 800 | 200 | | 800 | 200 | 800 | 200 | 800 |
| L-Cystine | 3 | 12 | 3 | | 12 | 3 | 12 | 3 | 12 |
| **Fiber** |  |  |  | |  |  |  |  |  |
| Cellulose, BW200 | 50 | 0 | 50 | | 0 | 50 | 0 | 50 | 0 |
| **Other components** |  |  |  | |  |  |  |  |  |
| Mineral Mix S10026 | 10 | 0 | 10 | | 0 | 10 | 0 | 10 | 0 |
| Dicalcium Phosphate | 13 | 0 | 13 | | 0 | 13 | 0 | 13 | 0 |
| Calcium Carbonate | 5.5 | 0 | 5.5 | | 0 | 5.5 | 0 | 5.5 | 0 |
| Potassium Citrate, 1 H2O | 16.5 | 0 | 16.5 | | 0 | 16.5 | 0 | 16.5 | 0 |
| Vitamin Mix V10001 | 10 | 40 | 10 | | 40 | 10 | 40 | 10 | 40 |
| Choline Bitartrate | 2 | 0 | 2 | | 0 | 2 | 0 | 2 | 0 |
|  | **gm** | **cal** | **gm** | | **cal** | **gm** | **cal** | **gm** | **cal** |
| **Total** | 1055.05 | 4057 | 773.85 | | 4057 | 904.05 | 4056 | 789.85 | 4057 |
| kcal/gm |  | 3.8 |  | | 5.2 |  | 4.5 |  | 5.1 |

**Supplemental Table 2. Sequence of qPCR primers used in this study**

| **Gene Name** |  | **NCBI Ref Seq acc#** | **Sense** | **Antisense** | **Product size (bp)** |
| --- | --- | --- | --- | --- | --- |
| ***Ppia*** | Peptidylprolyl isomerase A | NM_008907.1 | TGGTCTTTGGGAAGGTGAAAG | TGTCCACAGTCGGAAATGGT | 109 |
| ***Bactin*** | Beta actin | NM_007393.3 | CTGGGACGACATGGAGAAGA | ACCAGAGGCATACAGGGACA | 205 |
| ***Pparg*** | Peroxisome proliferator-activated receptor g | NM_001127330.1 | AGACCACTCGCATTCCTTTG | CCTGTTGTAGAGCTGGGTCTTT | 214 |
| ***Cidec*** | Cell Death Inducing DFFA Like Effector C | NM_178373.3 | AAGATGGCACAATCGTGGAG | TTAGTTGGCTTCTGGGAAAGG | 151 |
| ***Cidea*** | Cell Death Inducing DFFA Like Effector A | NM_007702.2 | GCAGCCTGCAGGAACTTATC | TCATGAAATGCGTGTTGTCC | 144 |
| ***Tnfa*** | Tumor necrosis factor alpha | NM_013693.2 | TAGCCCACGTCGTAGCAAAC | TGTGGGTGAGGAGCACGTA | 195 |
| ***Ccl2*** | Monocyte Chemoattractant Protein-1 | NM_011333.3 | CAGCAGGTGTCCCAAAGAAG | TGAGGTGGTTGTGGAAAAGG | 241 |
| ***Trem2*** | Triggering receptor expressed on myeloid cells 2 | NM_031254.3 | GGAACCGTCACCATCACTCT | CTTGATTCCTGGAGGTGCTGT | 220 |
| ***Col1a1*** | Collagen 1a1 | NM_007742 | TCAGAGGCGAAGGCAACA | AATGTCCAAGGGAGCCACA | 149 |
| ***Timp1*** | TIMP metallopeptidase inhibitor 1 | NM_001294280.2 | CCAGAACCGCAGTGAAGAG | CTCCAGTTTGCAAGGGATAGA | 193 |
| ***Mat1a*** | Methionine adenosyltransferase I, alpha | NM_133653.3 | TCTGCCAAAGATCTGCCTGT | TATCTGGATGCCCCTCTCCT | 207 |
| ***Gnmt*** | Glycine N-methyltransferase | NM_010321.1 | CAGAGTACAAGGCGTGGTTG | CTTTGTCCAGCGTCAACCAG | 243 |
| ***Pemt*** | Phosphatidylethanolamine N-methyltransferase | NM_001290011.1 | TAGCGAGATGGGAGCAGAGA | CTGGACAGCACAAACACGAA | 226 |
| ***Ahcy*** | S-adenosylhomocysteine hydrolase | NM_016661.3 | AAACCAGGTGATGGCACAGA | CAGCTGGAAGGTGAAGGACA | 241 |
| ***Bhmt*** | Betaine-homocysteine methyltransferase | NM_016668.3 | GAATTCCCCTTTGGATTGGA | CTGATCCAGGGTTTGGTGTG | 236 |
| ***Cbs*** | Cystathionine beta-synthase | NM_144855.3 | CCCATCCTTGCTGAGTTTGT | TCCAGACTCGATCCTTGTCC | 225 |

**Supplemental Figure 1.** Histological scores of steatosis (A), hepatocyte ballooning (B), and lobular inflammation (C) of hematoxylin & eosin (H&E)-stained liver section of mice in cohort #1 (G, LF, HF, and HFCF-fed mice) and cohort #2 (H, LF, HF, and HFC+Fr-fed mice). Letters indicate significant differences between LF-fed and HF-fed control mice. Exclamation marks (!) indicate significant differences between HF-fed and HFCF-control mice in Cohort #1 or HF-fed and HFC+Fr-control mice in Cohort #2. Plus signs (+) indicate significant differences between HF-fed and HFCF-*Pparg*^ΔHep^ mice in Cohort #1 or HF-fed and HFC+Fr-*Pparg*^ΔHep^ (KO) mice in Cohort #2. Asterisks indicate significant differences between control and *Pparg*^ΔHep^ mice. * p<0.05; ++, ** p<0.01; !!!,+++, p<0.001; d, !!!!, ++++, p<0.0001. n= 4-5 mice/group in cohort #1 and 5 mice/group in cohort #2.

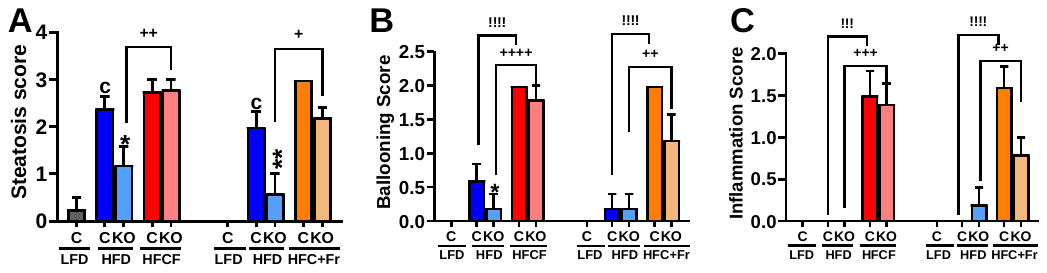

**Supplemental Figure 2.** Effect of sixteen weeks of HFC+Fr diet on the circadian changes for 96 hours in respiratory exchange ratio (RER, A), energy expenditure (B), food consumption (C), oxygen consumption (D), and carbon dioxide production (E) in male mice. Green lines represent data from LF-fed control mice, and red lines represent data from HFC+Fr-fed control mice. Effect of sixteen weeks of *Pparg*^ΔHep^on the circadian changes for 96 hours in respiratory exchange ratio (RER, F), energy expenditure (G), food consumption (H), oxygen consumption (I), and carbon dioxide production (J) in HFC+Fr-fed male mice. Red lines represent data of HFC+Fr-fed control mice, and blue lines represent data from HFC+Fr-*Pparg*^ΔHep^ control mice. n= 11-13 mice/group.

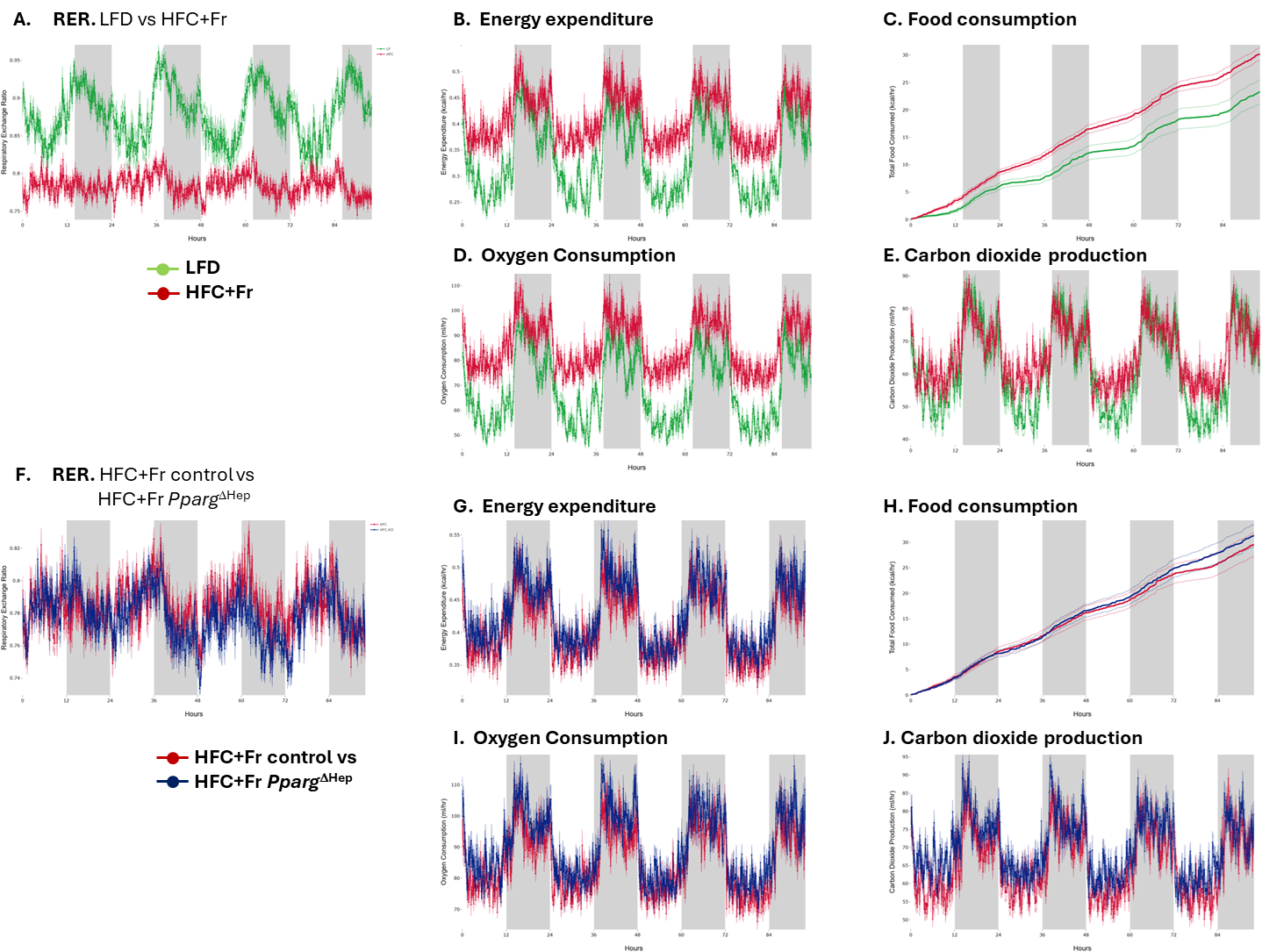

**Supplemental Figure 3.** ANCOVA of the mass and group independent effect of HFC+Fr diet or *Pparg*^ΔHep^ for sixteen weeks in male mice. The ANCOVA was performed for the variable energy expenditure, food consumption, oxygen consumption, and carbon dioxide production in the dark cycle (night, 2000h to 0600h) and light cycle (day, 0600h to 2000h). Asterisks show a significant effect independent of mass. **, p<0.01; ***, p<0.001. n= 11-13 mice/group.

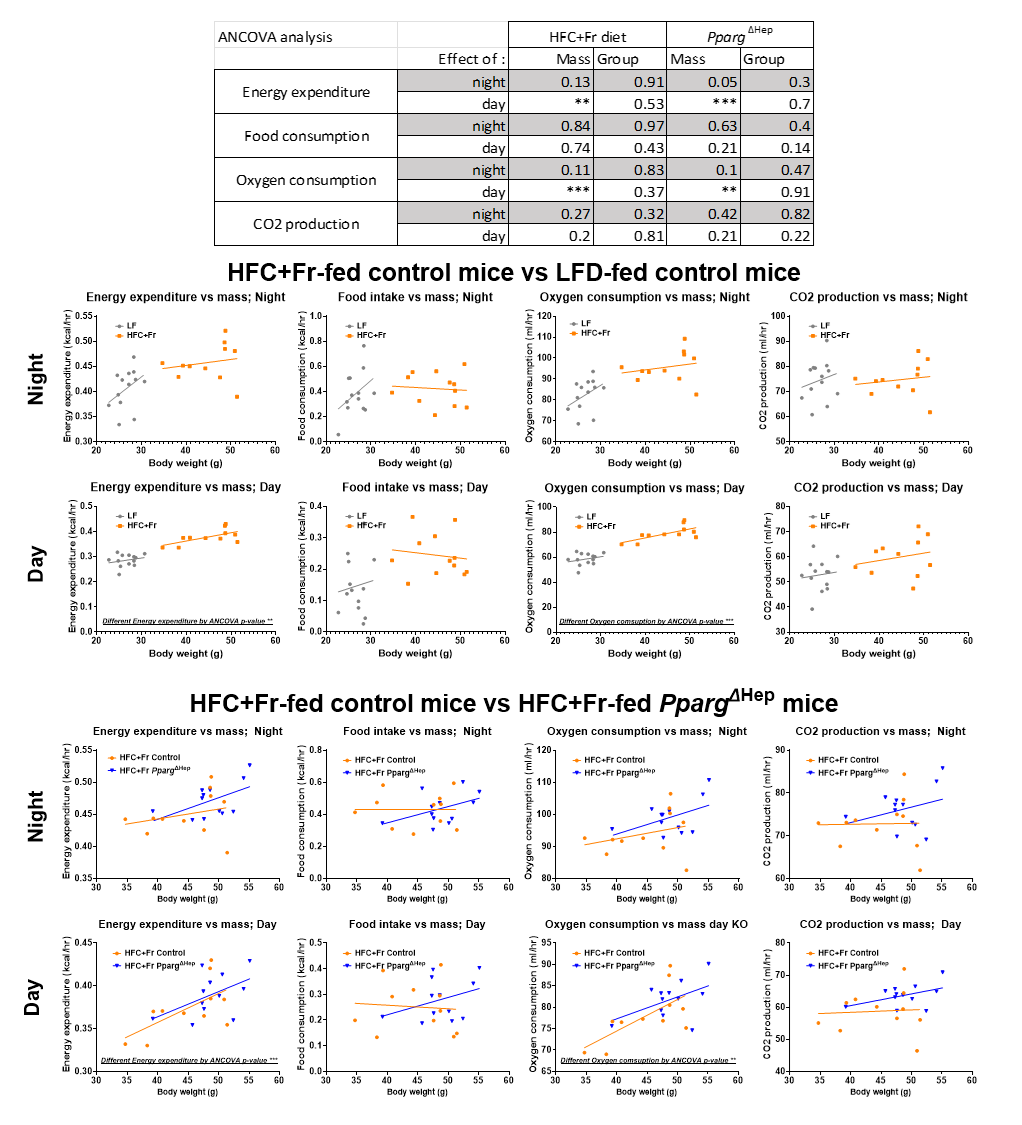

**Supplemental Figure 4.** Histological scores of steatosis (A), hepatocyte ballooning (B), and lobular inflammation (C) of hematoxylin & eosin (H&E)-stained liver section of male and female mice fed a HFC-Fr diet. Letters (a-d) indicate significant differences between LF-fed and HFC+Fr-fed control mice. Asterisks indicate significant differences between HFC+Fr-fed control and HFC+Fr-fed *Pparg*^ΔHep^ (KO) mice. a, * p<0.05; b, p<0.01; c, p<0.001; d, p<0.0001. n= 4-8 male mice/group and 5-7 female mice/group.

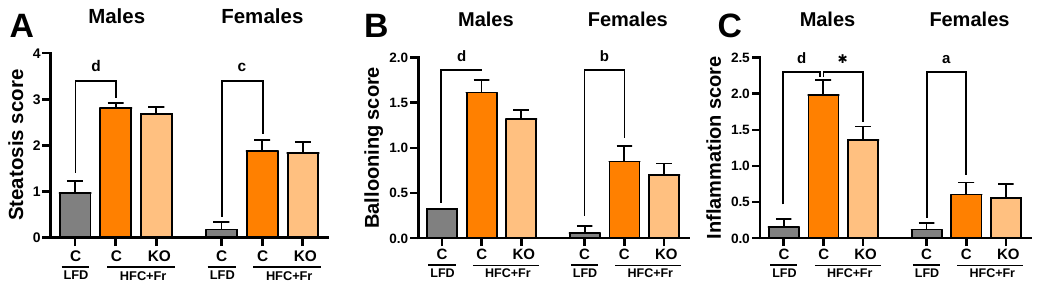

.
